# DSCAM-AS1 promotes tumor growth of breast cancer by reducing miR-204-5p and upregulating *RRM2*

**DOI:** 10.1101/400994

**Authors:** Wen-Hui Liang, Na Li, Zhi-Qing Yuan, Xin-Lai Qian, Zhi-Hui Wang

## Abstract

We intended to analyze the effects of DSCAM-AS1, miR-204-5p and *RRM2* on breast cancer (BC) cells growth. Microarray analysis and qRT-PCR were employed to determine DSCAM-AS1 and miR-204-5p expression in BC. Luciferase reporter assay and cell transfection assay were applied to examine the target relationship between DSCAM-AS1, miR-204-5p and MMR2. CCK-8 assay, transwell assay and flow cytometry were used to detect cell proliferation, invasion and apoptosis of breast cancer cells. The expression of DSCAM-AS1, miR-204-5p and MMR2 were confirmed by Western Blot. We also conducted *In vivo* assay to verify the effect of DSCAM-AS1 on tumor formation.DSCAM-AS1 was up-regulated, while miR-204-5p was down-regulated in BC tissues and cells. Meanwhile, DSCAM-AS1 directly targeted miR-204-5p. DSCAM-AS1 promoted the proliferation and invasion of BC cells and restrained cell apoptosis by reducing miR-204-5p and inhibiting miR-204-5p expression. *RRM2* was up-regulated in BC cells, and miR-204-5p inhibited *RRM2* expression by targeting *RRM2*. Overexpression of *RRM2* stimulated proliferation and cell invasion and impeded apoptosis of BC cells. *In vivo* experiments showed that knockdown of DSCAM-AS1 decreased the tumorigenesis of BC cells, increased the expression of miR-204-5p while inhibited *RRM2* expression.DSCAM-AS1 promoted proliferation and impaired apoptosis of BC cells by reducing miR-204-5p and enhancing *RRM2* expression. DSCAM-AS1/miR-204-5p/*RRM2* may serve as novel therapeutic targets for BC.

**Summary statement:** Microarray analysis and qRT-PCR were employed to determine DSCAM-AS1 and miR-204-5p expression in BC. DSCAM-AS1 promoted proliferation and impaired apoptosis of BC cells by reducing miR-204-5p and enhancing *RRM2* expression.

## Introduction

Breast cancer (BC) is a common malignancy around the world, and more than 370,000 women die due to BC worldwide each year (Wang et al., 2017). The main cause of BC morbidity and mortality is an incurable metastatic disease that is highly resistant to traditional therapies (Xu et al., 2017). Despite the advances in diagnosis and therapy, such as surgery, chemotherapy and radiation, there are high rates of recurrence and metastasis among patients with BC (Wang et al., 2015). Besides, continuous treatment with radiotherapy and chemotherapy leads to less effective destruction of cancer cells due to the acquired resistance (Vimalraj et al., 2013). Therefore, the molecular mechanisms underlying BC tumorigenesis need to be further investigated.

Long noncoding RNAs (lncRNAs) are defined as transcripts that have more than 200 nucleotides and lack protein-coding capacity (Xu et al., 2017). LncRNAs have been reported to involve in tumor progression and metastasis through regulating different levels biological processes, such as mRNA alternative splicing, protein activities, changes of protein localization, etc (Huarte, 2015; Xu et al., 2017). A growing number of literatures reported that lncRNAs are tightly linked to human cancer and involved in the control of availability for specific miRNAs (Bartonicek et al., 2016). In BC, a number of differentially expressed lncRNAs and their correlation with tumorigenic functions were reported (Xi et al., 2017; Yan et al., 2017; Zhou et al., 2017). For example, lncFOXO1 was notably down-regulated in BC and inhibited the growth of BC by increasing FOXO1 transcription (Xi et al., 2017). While, linc-ITGB1 was greatly up-regulated in BC which promoted cell metastasis (Yan et al., 2017).

DSCAM (Down Syndrome Cell Adhesion Molecule) antisense lncRNA (DSCAM-AS1), located on 21q22.2, is transcribed from the antisense strand of DSCAM belonging to the immunoglobulin superfamily of cell adhesion molecules (Zhao et al., 2014). Several literatures have reported that DSCAM-AS1 is associated with the progression of cancer cells (Miano et al., 2016; Niknafs et al., 2016; Zhao et al., 2014). Zhao *et al*. found that DSCAM-AS1 was overexpressed in lung adenocarcinoma and might interact with their host genes to accomplish the function of different subtypes of lung cancer (Zhao et al., 2014). Miano *et al*. discovered that overexpression of DSCAM-AS1 promoted the growth of a luminal breast cancer cell line (Miano et al., 2016). Of note, Yashar S et al. showed that DSCAM-AS1 could interact with hnRNPL to mediate tumor progression (Niknafs et al., 2016). However, the whole story of how DSCAM-AS1-mediated luminal breast cancer growth and progression largely remains unkown. Therefore, a further understanding of DSCAM-AS1-regulated targets and networks is needed.

MicroRNAs (miRNAs) are small noncoding RNA molecules of 19-25 nucleotides and can act as post-transcriptional regulators by binding to complementary sequences of target mRNAs (Zeng et al., 2016). MiRNAs were involved in cellular processes including cell proliferation, differentiation and apoptosis (Li et al., 2014; Liu and Li, 2015; Wang et al., 2017; Zeng et al., 2016). A host of evidences have revealed that dysregulation of miRNAs is associated with BC tumorigenesis and metastasis. MiR-204-5p is a type of microRNA, and previously studies have reported that miR-204-5p was down-regulated in several human cancers and might act as a tumor inhibitor (Luo et al., 2017). For example, low levels of miR-204-5p expression were confirmed in hepatocellular carcinoma (Luo et al., 2017), oral squamous cell carcinoma (Wang et al., 2016), gastric cancer (Zhang et al., 2015), papillary thyroid carcinoma (Liu et al., 2015) and glioma (Xia et al., 2015). And miR-204-5p has been confirmed to associate with BC cells, and overexpression of miR-204-5p might suppress cell migration and invasion of BC (Zeng et al., 2016). Besides, Wang *et al.* reported that miR-204 interacted with lncRNA MALAT1 and promoted BC cell invasion through inducing epithelial–mesenchymal transition (Wang et al., 2017), in microscopic view, Shen et al. showed tha miR-204 suppressed the breast cancer cells via directly targeting FOXA1 (Shen et al., 2017); Flores et al. reported miR-204 controlled the angiogenesis in breast cancer regulating ANGPT and TGFBR2 (Flores-Perez et al., 2016); while Imam et al. showed loss of miR-204 promoted breast cancer migration and invasion via activating AKT/mTOR/Rac1 signaling and actin reorganization (Imam et al., 2012). Yet no study focused on the correlation between miR-204-5p and DSCAM-AS1.

Ribonucleotide reductase M2 (*RRM2*) is the catalytic subunit of ribonucleotide reductase which is an essential enzyme involved in DNA synthesis, and can regulate its enzymatic activity (Iwamoto et al., 2015). Several publications have reported that *RRM2* was overexpressed in diverse cancer cells (Zhong et al., 2016). Dysregulation of *RRM2* in BC has been previously investigated in some literatures. For instance, *RRM2* was up-regulated in BC cells and could act as a critical marker for aggressive BC (Lu et al., 2012). Shah *et al.* found that inhibition of *RRM2* suppressed *in vivo* tumor growth and decreased cell migratory and invasive in BC (Putluri et al., 2014). Several studies indicated the role of *RRM2* as a target of miRNA in human cancers. The results of our experiments clarified the regulation mechanism between *RRM2* and miR-204-5p.

In current study, we measured the expression level of DSCAM-AS1 and miR-204-5p in BC cells. Subsequently, we employed dual-luciferase reporter system to verify the correlation of DSCAM-AS1 and miR-204-5p. Afterwards, the effects of DSCAM-AS1 on BC proliferation, growth metastasis and apoptosis were measured by a series of *in vitro* experiments including cell apoptosis assay, CCK-8 assay and transwell assay and *in vivo* experiments. In addition, we also explored the impacts of *RRM2* on BC cell activities and tumor growth. These results highlighted that DSCAM-AS1 may be a promising therapeutic strategy for treatment of human BC.

## Materials and methods

### Tissue specimens

BC tissues and adjacent tissues were collected between January 2014 and August 2015 from 40 patients of the Affiliated Center Hospital, Xinxiang Medical University. All specimens were gathered from women aged 44-71 (average age: 62.3). Patients did not receive any chemotherapy or radiation therapy before surgery. All tissue samples were independently checked by three pathologists or doctors before preserving in liquid nitrogen at −80°C or transferring to microcentrifuge tubes containing TRIzol reagent for RNA extraction. The relevant characteristics of these 40 patients are described in Table 1. All experiments were performed with the informed consent of the patients and under the approval of the Affiliated Center Hospital, Xinxiang Medical University.

**Table 1.**
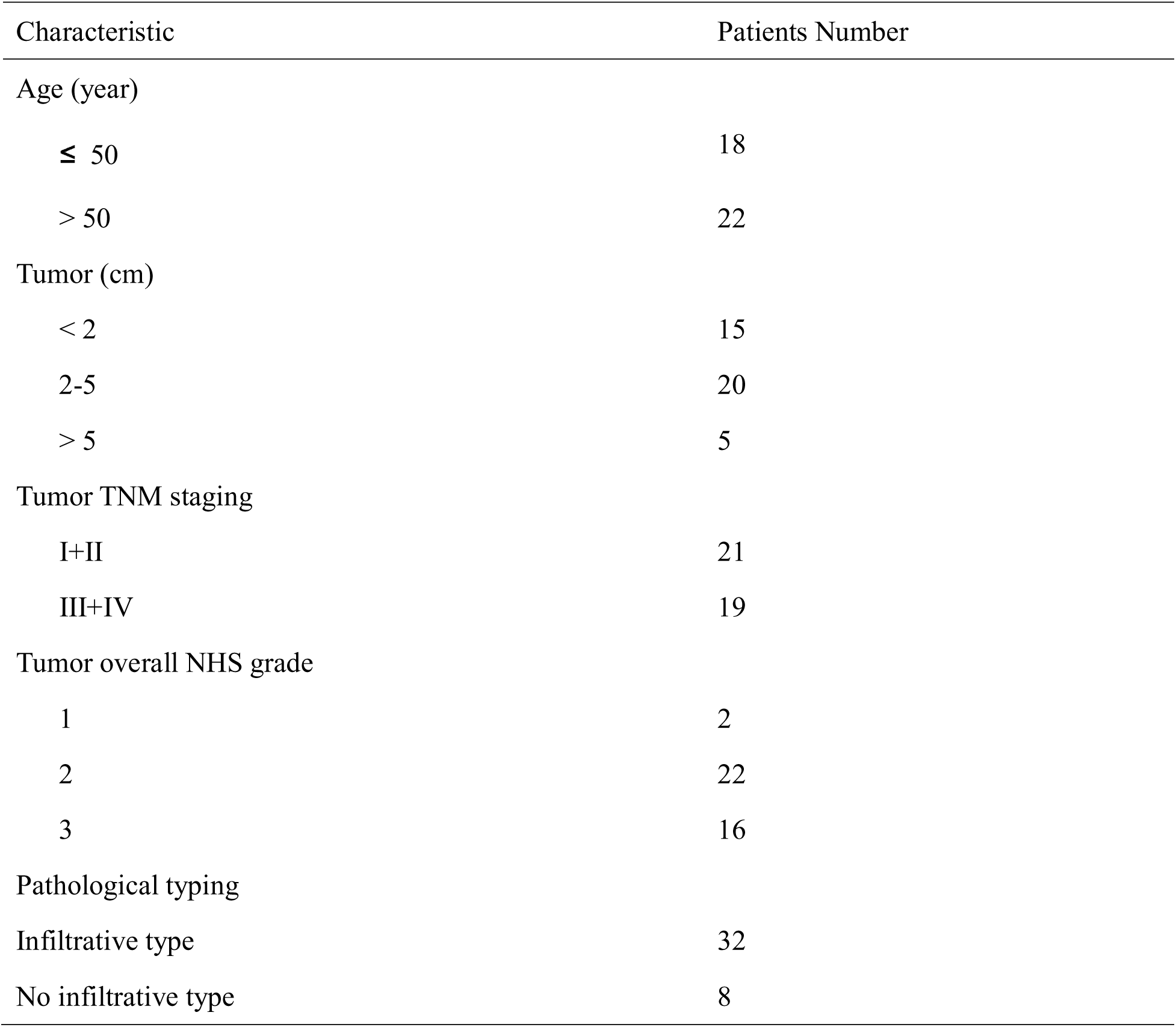
Clinical and pathologic characteristics of study subjects

| Characteristic | Patients Number |
| --- | --- |
| Age (year) |  |
| $\leq 50$ | 18 |
| $> 50$ | 22 |
| Tumor (cm) |  |
| $< 2$ | 15 |
| 2-5 | 20 |
| $> 5$ | 5 |
| Tumor TNM staging |  |
| I+II | 21 |
| III+IV | 19 |
| Tumor overall NHS grade |  |
| 1 | 2 |
| 2 | 22 |
| 3 | 16 |
| Pathological typing |  |
| Infiltrative type | 32 |
| No infiltrative type | 8 |

### Cell culture

Human breast cancer cell line HCC1937 and human embryonic kidney cell line HEK293 were purchased from Chinese Academy of Sciences cell bank (Shanghai, China). HCC1937 cells were cultured in the RPMI-1640 medium (GIBCO, Grand Island, NY, USA) plus 10% (v/v) fetal bovine serum (FBS), 1.5g/L of NaHCO_3_, 2.5g/L glucose, 0.11 g/L sodium pyruvate and 1%(v/v) penicillin-streptomycin. HEK293 cells were maintained in DMEM medium supplemented with 10%(v/v) FBS AND 1%(v/v) penicillin-streptomycin.

### Microarray analysis

The breast tumor tissue data was collected from TCGA database. The ‘DESeq2’ R-package was utilized to normalize, identify and visualize differentially expressed miRNAs and lncRNAs (log_2_|Fold Change|>1 and *P.adjust*<0.05). The ‘pheatmap’ R-package was used to generate heatmaps of top 17 differentially expressed RNAs via pheatmap function. After normalization, we screened out differentially expressed lncRNAs from 48 luminal, 21 TNBC, and 22 Her2 positive breast cancer tissues compared to adjacent normal tissues, and differentially expressed miRNAs in 43 luminal, 19 TNBC, and 22 Her2 positive breast cancer tissues compared to adjacent normal tissues.

### Cell transfection

Overexpression of DSCAM-AS1 (isoform 2), the predominant isoform, was obtained by PCR and cloned into pLenti6.1 vector (Invitrogen, CA, USA), and shDSCAM-AS1(shRNA) was purchased from GenePharma (Shanghai, China). For in vitro assays, transient transfection in HCC1937 cells was performed using Lipofectamine 3000 (Invitrogen, CA, USA) for pLenti6.1-DSCAM-AS1 and pLenti6.1-shDSCAM-AS1, MiR-204-5p mimics (GenePharma, Shanghai, China), and miR-NC(non-targeting control) (GenePharma, Shanghai, China). Cells were divided into different groups by two ways. The first group settings were as follows: cells transfected with pLenti6.1 and miR-NC served as NC group; cells transfected with pLenti6.1 and miR-204-5p was regarded as miR-204-5p group; cells transfected with pLenti6.1-DSCAM-AS1and miR-NC served as DSCAM-AS1 group; cells transfected with pLenti6.1-DSCAM-AS1 and miR-204-5p was regarded as DSCAM-AS1+miR-204-5p group. The second group settings were as follows: cells transfected with pLenti6.1 and miR-NC served as NC group; cells transfected with pLenti6.1 and miR-204-5p was regarded as miR-204-5p group; cells transfected with pcDNA3.1-*RRM2* and miR-NC served as *RRM2* group; cells transfected with pcDNA3.1-*RRM2* and miR-204-5p was regarded as *RRM2*+miR-204-5p group. Transfection was performed by Lipofectamine 3000 (Invitrogen, CA, USA) following the manufacturer’s protocols. All fragments and inserts were confirmed by sequencing.

### QRT-PCR

TRIzol Reagent (Invitrogen, Carlsbad, CA, USA) and NanoDrop 2000 (Thermo Fisher Scientific Inc., USA) were used to extract and quantify total RNA, respectively. Reverse transcription was performed with 200 ng total RNA by using PrimeScript High Fidelity RT-PCR Kit (Takara, Japan). QRT-PCR was performed using PowerUp™ SYBR™ Green Master Mix (Thermo-Fisher) with a QuantStudio™ 3 Real-Time PCR System(Thermo-Fisher) according to the manufacturer’s instructions. U6 and GADPH were used as internal references. 2^-ΔΔCt^ method was employed to quantify relative mRNA expression values. The specific primers used are presented in Table 2.

**Table 2.**
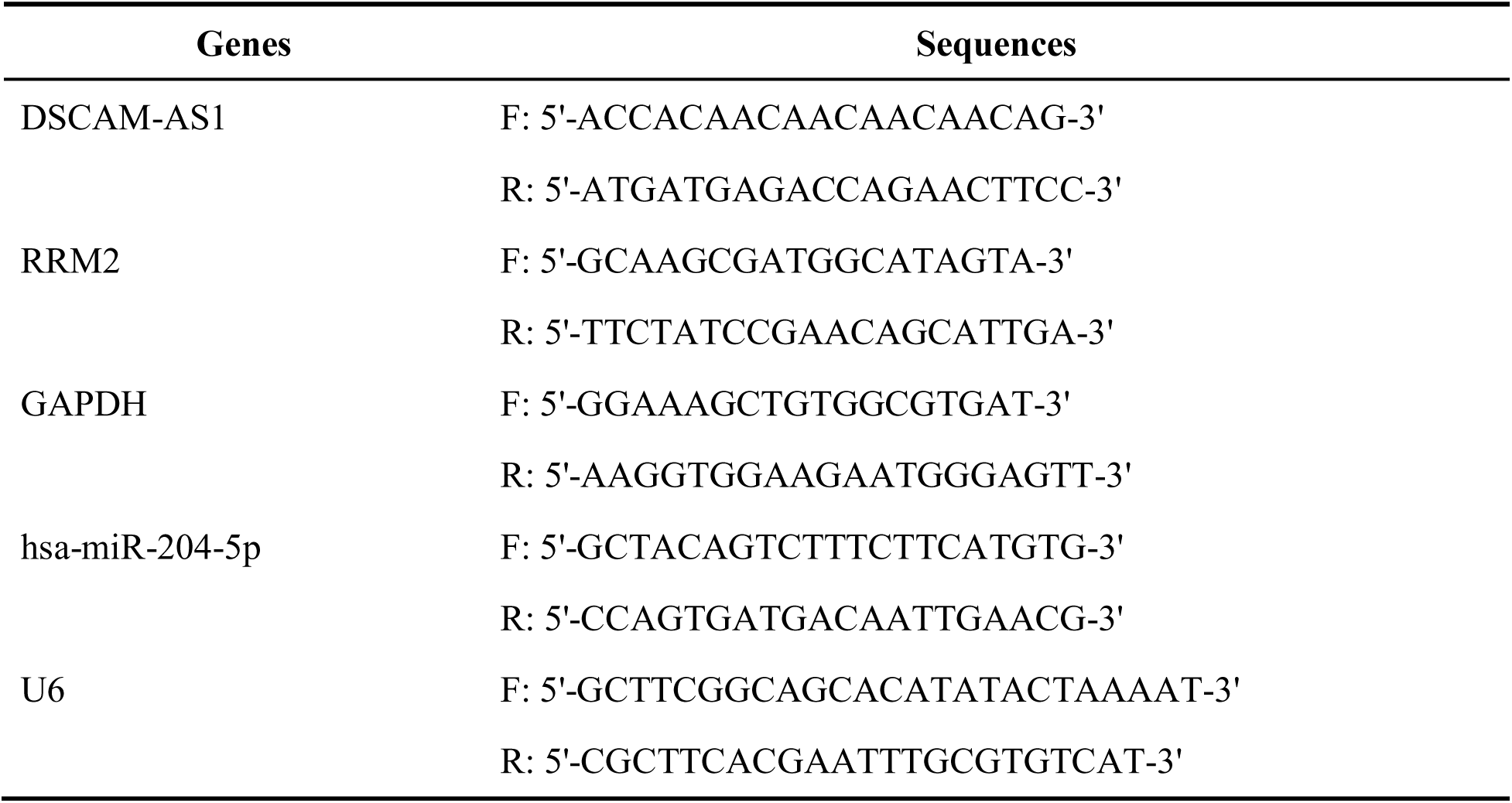
Primer Sequences for qRT-PCR

### Western blot

Protein samples were lysed in RIPA buffer (Beyotime, Shanghai, China) with a cocktail of protease inhibitor (Roche, Basel, Switzerland). After centrifugation, supernatant protein concentration was measured by Bradford Reagent (Bio-Rad, CA, USA). After eluted with SDS-loading buffer, proteins were electrophoresed in 10% SDS-polyacrylamide gel electrophoresis (SDS-PAGE) gel, transferred to nitrocellulose membranes (Millipore, USA) and blocked. Primary antibodies (anti-*RRM2*, ab172476, 1:2000; anti-GAPDH, ab181603, 1:10000, Abcam, Cambridge, MA, USA) were added. After extensively washed with TBST, secondary anti-body was added (Goat anti-Rabbit IgG H&L (HRP), ab6721, 1:2000, Abcam, Cambridge, MA, USA) for 1.5 h. Signal detection was carried out by immunoblotting with an ECL system (Life Technology, USA). For relative quantification, we normalized GADPH expression level (as gray level) to 1, and compared the expression levels (gray level) of different experimental groups via ImageJ software.

### Flow cytometry

Cell apoptosis was assessed by Annexin V-PE Apoptosis Detection Kit (Beyotime, Shanghai, China) following the manufacturer’s protocols. 48 hours after transfection, cells were washed re-suspended in binding buffer containing propidium iodide (PI) and annexin V-FITC. Stained cells were analyzed by BD Accuri C6 flow cytometer (BD, USA) using Cell Quest Pro software (BD, USA).

### CCK-8 assay

8×10^3^ HCC1937 cells per well were incubated into 96 well and cell vitality was assessed by Cell Counting Kit-8 (Biotechwell, Shanghai, China) at 0, 24, 48, 72 and 96 h according to the manufacturer’s instructions. Absorbance was recorded at 450 nm with a SpectraMax i3x Multi-Mode Detection Platform (Molecular Devices, USA).

### Transwell assay

Cells were suspended in serum-free medium and placed into the upper chamber with coated Matrigel of a 24-well chamber (Sigma-Aldrich, MO, USA). Six-hundred-microliter media containing 10% FBS were added to the lower chamber overnight. Migrated cells were fixed with 4% paraformaldehyde and stained with crystal violet. Cells were rinsed and counted from random fields (×100 magnification). Each experiment was conducted triplicate.

### Luciferase reporter assays

Twenty-four hours before transfection HEK293 cells were seeded onto a 24-well plate (1×10^5^/well), then the cells were co-transfected with pCMV-Renilla (Promega, WI, USA) and firefly luciferase reporter plasmids pGL3 (Promega, WI, USA) containing negative control (NC), or 3’UTR of miR-204-5p downstream of the *Firefly* luciferase gene, along with pLenti6.1-DSCAM-AS1, pLenti6.1-DSCAM-AS1 mut#1(region 406-427 mutated), or pLenti6.1-DSCAM-AS1 mut#2(region 685-706 mutated), and pcDNA3.1(+)-RRM2 or pcDNA3.1(+)-RRM2 mut#(region 1949-1956 mutated) using Lipofectamine 3000 (Invitrogen, CA, USA) according to the manufacturer’s recommendations. The mutants were generated by PCR and the sequence was shown in Fig2A-B, which were confirmed by DNA sequencing. Forty-eight hours after transfection, luciferase activities were performed using Dual-luciferase Reporter Assay Kit (Promega, WI, USA). Then the firefly and renilla luciferase activities were measure by a Berthold (Bundoora, VIC, Australia) luminometer.

### Animal experiment

For in vivo trials, HEK293T were transduced in the presence of 8ug/ml polybrene and grown in culture medium containing 2ug/ml puromycin to generate pLenti6.1-DSCAM-AS1 and pLenti6.1-shDSCAM-AS1 lentiviral particles. HCC1937 cells were transduced by these lentiviral particles. Then, cells were cultured in the medium with 3ug/ml blasticidin for stable cell line selection before subcutaneously injected into the right flank of female nude mice(6-8 weeks old). The tumor sizes were measured every 5 d using caliper. The volume of tumor was estimated using the fomula V=(π/6)(W^2*^L), where W=width and L=length of the tumor. 30 days post-injection, mice were sacrificed and the size of tumor was used as the endpoint of reading. All animal studies were approved by the Affiliated Center Hospital, Xinxiang Medical University and in accordance with the norms of The Affiliated Center Hospital, Xinxiang Medical University. The data are presented as the mean volume X±S.E.

### Statistical analysis

All experiments were repeated at least three times. All statistical analyses were performed using GraphPad Prism 6.0 software (GraphPad Software Inc., USA). The measured parameters are presented as the mean ± SD. One-way analysis of variance (ANOVA) followed by Tukey’s test were used to compare the differences among groups. Statistical significance is presented in figures by *, *P*<0.05, **, *P*<0.01.

## Results

### Identification of DSCAM-AS1 is an up-regulated lncRNA in breast tumor

We initially focused on investigating the most differentially expressed lncRNAs and miRNAs in breast cancer tissue compared to benign adjacent tissue from 1097 tissue samples. All differentially expressed lncRNAs and miRNAs with a filtration standard of log_2_|Fold Change|>1 and *P.adjust*<0.05 were demonstrated in a volcano map (Figure 1A-B). Among all top differentially expressed lncRNAs, we noticed that DSCAM-AS1 plays a crucial role in breast tumor progression and migration. Because DSCAM-AS1 was reported to highly expressed in luminal breast cancer specifically (Miano et al., 2016; Niknafs et al., 2016), we decided to visualize top differentially expressed lncRNAs and miRNAs in luminal breast cancer tissues. A group of 18 lncRNAs with 10 down-regulated and 7 up-regulated and another group of 17 miRNAs with 8 down-regulated and 9 up-regulated in TCGA BC tumor tissues compared to TCGA-adjacent tissues were shown in heat map (Figure 1C-D). Total identified differentially expressed lncRNAs and miRNAs were listed in supplementary Table.1-2. Consistent with previous reports, our data supported that DSCAM-AS1 was highly expressed in luminal breast cancer tissues in comparison to adjacent normal tissues. We also found others researches have indicated that miR-204-5p is a breast tumor suppressor gene, which is also in line with our analysis. Thus, we were eager to further validate our bioinformatics analysis, we employed qRT-PCR to detect DSCAM-AS1 levels in BC tissue and normal adjacent tissue samples of forty hospital patients. The results indicated that in comparison with normal adjacent tissues, DSCAM-AS1 was up-regulated (Figure 1E, *P*<0.05) while miR-204-5p was down-regulated (Figure 1F, *P*<0.05) in BC tissue samples.

**Fig. 1.**
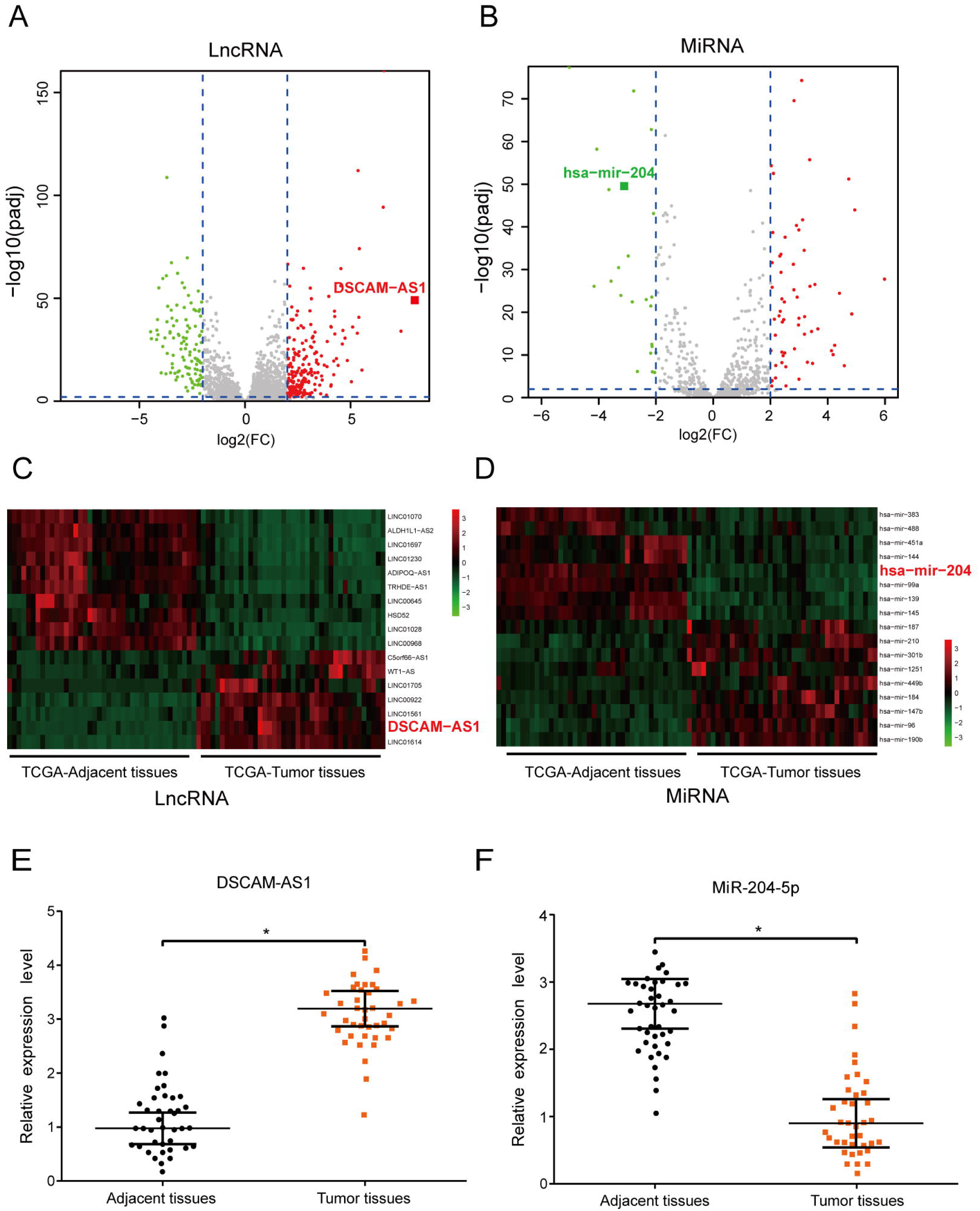
DSCAM-AS1 was up-regulated and miR-204-5p was down-regulated in BC tissues. (A-B) The volcano map showed differentially expressed lncRNAs and miRNAs screened by log_2_|Fold Change|> 2, *P.adjust* <0.05. (C) Heatmap indicated that 7 up-regulated and 10 down-regulated mRNAs in luminal BC tumor tissues. (D) Heat map indicated 9 up-regulated and 8 down-regulated miRNAs in luminal BC tumor tissues. (E) qRT-PCR revealed that DSCAM-AS1 expression in tumors was much higher than that in adjacent tissues. (F) qRT-PCR showed that miR-204-5p expression in tumors was significantly lower than that in adjacent tissues. ^**^ *P*<0.05,

### DSCAM-AS1 target on miR-204-5p

Then, we were wondering the target candidates of DSCAM-AS1 and miR-204-5p. Interestingly, via Targetscan and Miranda database, we predicted two potential target sites (Region 406-427 and Region 685-706) between DSCAM-AS1 and 3’UTR of miR-204-5p. Thus, we constructed the luciferase reporter and performed the trials as mentioned in the Material&Methods section, the results verified our prediction and confirmed that DSCAM-AS1 could regulate the 3’UTR of miR-204-5p (Figure 2A-B). The results of qRT-PCR showed that cells were successfully transfected with DSCAM-AS1, and the content of miR-204-5p was impaired when overexpressing DSCAM-AS1 (Figure 2C-D). In conclusion, DSCAM-AS1 could regulate on 3’UTR of miR-204-5p.

**Fig. 2.**
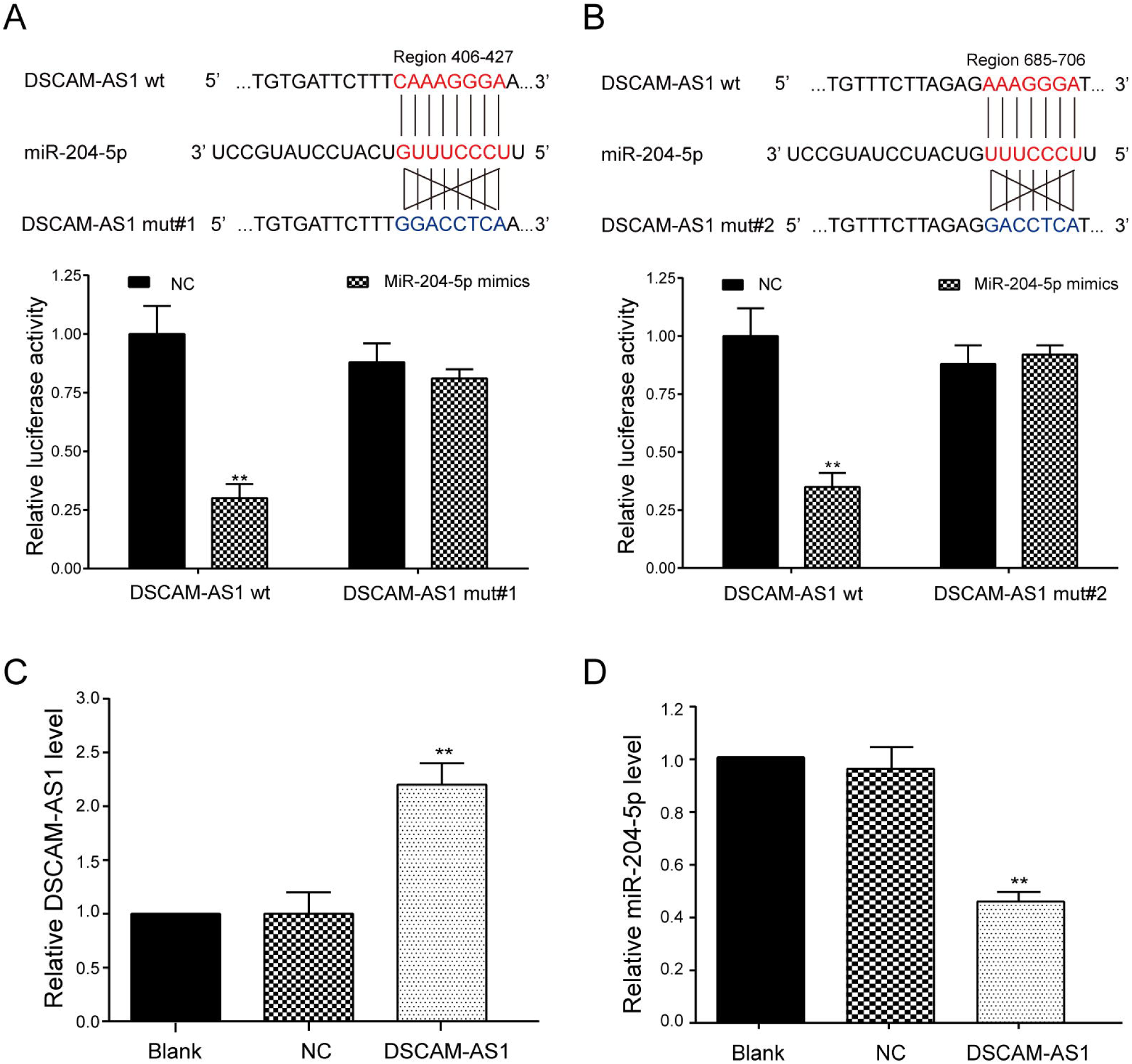
DSCAM-AS1 could directly target on miR-204-5p. (A) The targeted sites of miR-204-5p (Region 406-427 and Region 685-706) and experimental validation of them by dual luciferase reporter assays. ^**^ *P*<0.01, compared with NC+DSCAM-AS1 wt group. (B) The targeted site of DSCAM-AS1 (Region 685-706) and miR-204-5p and experimental validation of dual-luciferase reporter assays. ^**^ *P*<0.01, compared with NC+DSCAM-AS1 wt group. (C) QRT-PCR revealed that DSCAM-AS1 expression increased in HCC1937 cells transfected with DSCAM-AS1. ^**^ *P*<0.01, compared with NC group. (D) QRT-PCR revealed that miR-204-5p expression decreased in HCC1937 cells transfected with DSCAM-AS1. ^**^ *P*<0.01, compared with NC group.

### DSCAM-AS1 promoted BC cells proliferation and metastasis and impeded cell apoptosis by targeting miR-204-5p

The experiments were performed in four groups: NC group, miR-204-5p group, DSCAM-AS1 group and miR-204-5p+DSCAM-AS1 group. CCK-8 results showed that DSCAM-AS1 promoted the proliferation of BC cells, which was inhibited by miR-204-5p (Figure 3A, *P*<0.01). Transwell assay results demonstrated that miR-204-5p had opposite effect on cell migration, while DSCAM-AS1 enhanced BC cells migration, and co-transfection of DSCAM-AS1 and miR-204-5p had remarkably reversed the DSCAM-AS1 induced BC cells migration and invasion (Figure 3B-D, *P*<0.01). Flow cytometry analysis revealed that BC cell apoptosis was impaired by DSCAM-AS1 and stimulated by miR-204-5p, and co-transfection of DSCAM-AS1 and miR-204-5p had remarkably enhanced the BC cells apoptosis inhibited by DSCAM-AS1 (Figure 3E, *P*<0.01). Therefore, miR-204-5p served as tumor inhibitor by impeding the acceleration of DSCAM-AS1 on BC cells progression. Taken together, these panels demonstrated that DSCAM-AS1 has an inhibitory effect on miR-204-5p to promote BC cells proliferation and metastasis.

**Fig. 3.**
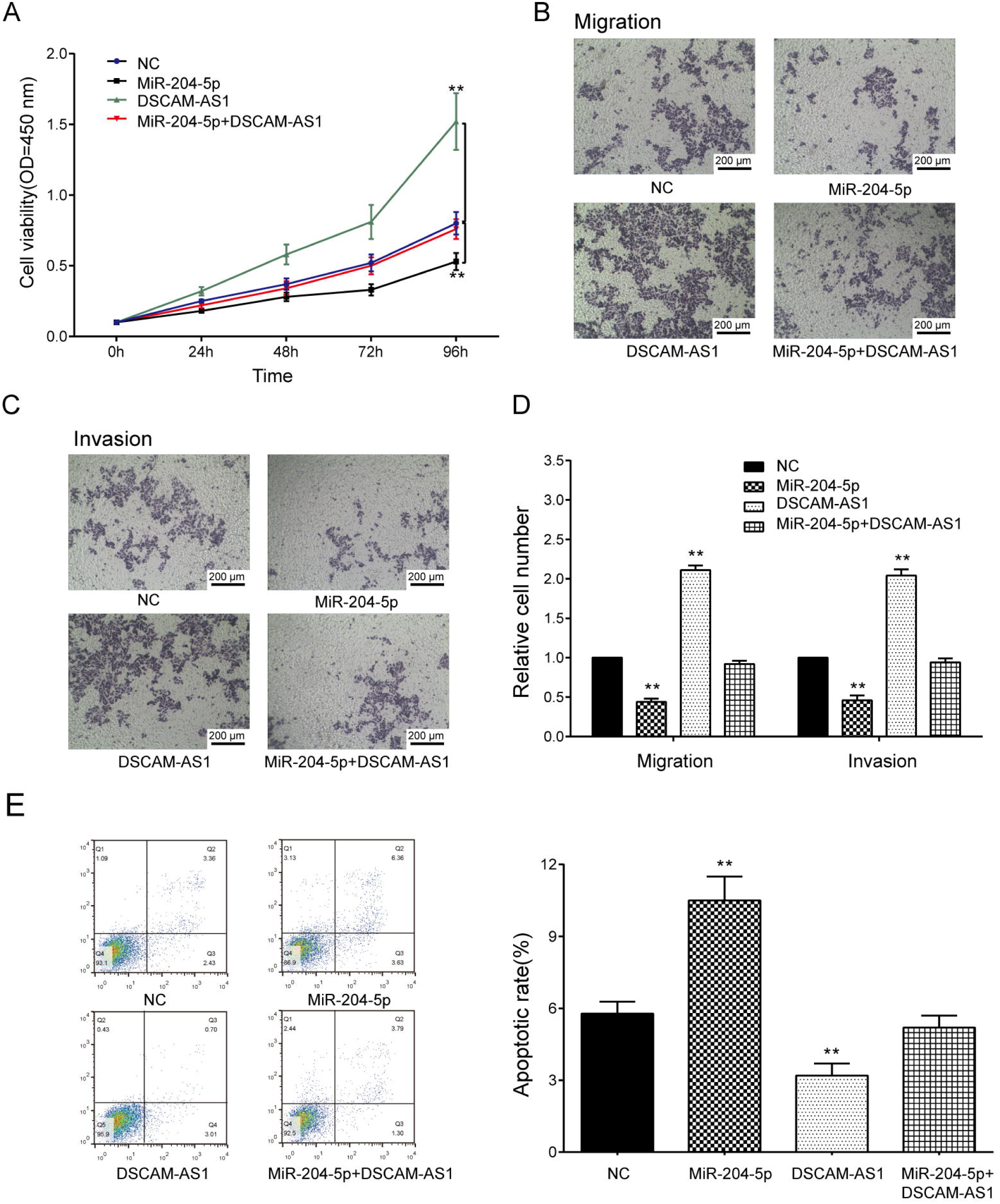
Effect of DSCAM-AS1 and miR-204-5p on BC cells activities. (A) Cell vitality of HCC1937 cells transfected with DSCAM-AS1 was higher than that of cells in NC group, while that in miR-204-5p group was lower than that in NC group detected by CCK-8 assay. (B-D) Transwell assays revealed that compared with NC group, more migrating or more invasive cells were found in DSCAM-AS1 group while less metastasis cancer cells were observed in miR-204-5p group. (E) Flow cytometry showed that apoptosis rate in DSCAM-AS1 group was lower than that in NC group, while that in miR-204-5p group was much higher than that in NC group. ^**^ *P*<0.01, compared with NC group.

### MiR-204-5p could act on *RRM2*

We hypothesized that *RRM2* was a downstream of miR-204-5p and the target sites between them were predicted by using Targetscan and Miranda database. We mutated Region 1949-1956 of RRM2 as mentioned in the left panel Figure 4A. As verified by dual luciferase reporter assay, we confirmed that RRM2 is a target of miR-204-5p (right panel, Figure 4A, P<0.01). Meanwhile, we used QRT-PCR and western blot to separately detect the mRNA and protein expression of RRM2 in BC tissue and adjacent tissue samples. As shown in Figure 4B and 4C, we verified higher expression of *RRM2* in BC tissues (Figure 4B-C, both *P*<0.05). In order to investigate the regulation of miR-204-5p on RRM2, we transfected HCC1937 cells with three different constructs or constructs combinations, the qRT-PCR results revealed that the expression of *RRM2* were significantly inhibited by miR-204-5p (Figure 4D-E, *P*<0.01).

**Fig. 4.**
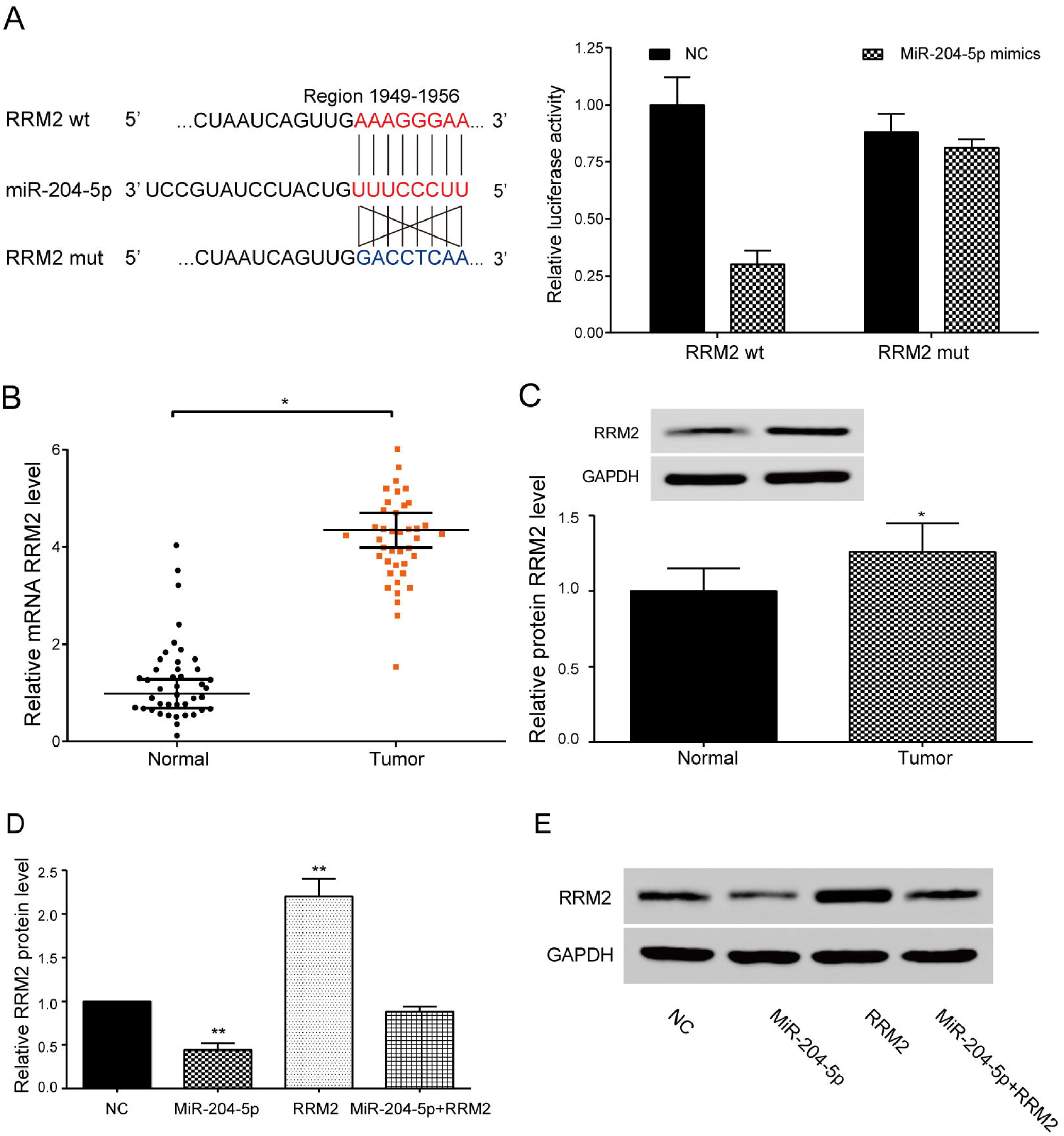
MiR-204-5p could directly target on RRM2. (A) The targeted site of miR-204-5p and RRM2 and experimental validation of it by dual luciferase reporter assays. ^**^ *P*<0.01, compared with NC+ RRM2 wt group. (B) QRT-PCR revealed that RRM2 expression was up-regulated in tumor tissues. ^*^ *P*<0.05, compared with normal tissues. (C) Western blot showed that RRM2 protein expression in tumor tissues was higher than that in normal tissues. (D) QRT-PCR revealed that RRM2 expression in miR-204-5p group was significantly lower than that in NC group. ^**^ *P*<0.01, compared with NC group. (E) Western blot showed that RRM2 protein expression decreased in HCC1937 cells when co-transfected with miR-204-5p.

### *RRM2* promoted BC cells reproduction and metastasis and impeded cell apoptosis which was reversely changed by miR-204-5p

Experimental groups were performed as follows: NC group, miR-204-5p group, *RRM2* group and miR-204-5p+*RRM2* group. The CCK-8 results revealed that the miR-204-5p effectively reduced BC cells viability while *RRM2* increased it, and in miR-204-5p+*RRM2* group, miR-204-5p largely suppressed the BC cells survival rate which was largely increased by RRM2. (Figure 5A, *P*<0.01). We used transwell assays to detect RRM2’s function on BC cell migration and invasion. As shown in Figure 5B-5D, miR-204-5p effectively inhibited the migration and invasion of BC cells, whereas RRM2 significantly enhanced these abilities of BC cells, and in co-transfection group of RRM2 and miR-204-5p, miR-204-5p had largely inhibited the migration and invasion of BC cells which was increased by RRM2. Next, we applied cell apoptosis assay and Flow cytometry technique to confirm RRM2’s function on BC cell apoptosis. The results indicated that *RRM2* impeded cancer cell apoptosis, while the inhibitory effect of *RRM2* was reversely changed by the participation of miR-204-5p (Figure 5E, *P*<0.01). Together, these results revealed that miR-204-5p may have an inhibitory effect on RRM2 to suppress BC cells growth and progression.

**Fig. 5.**
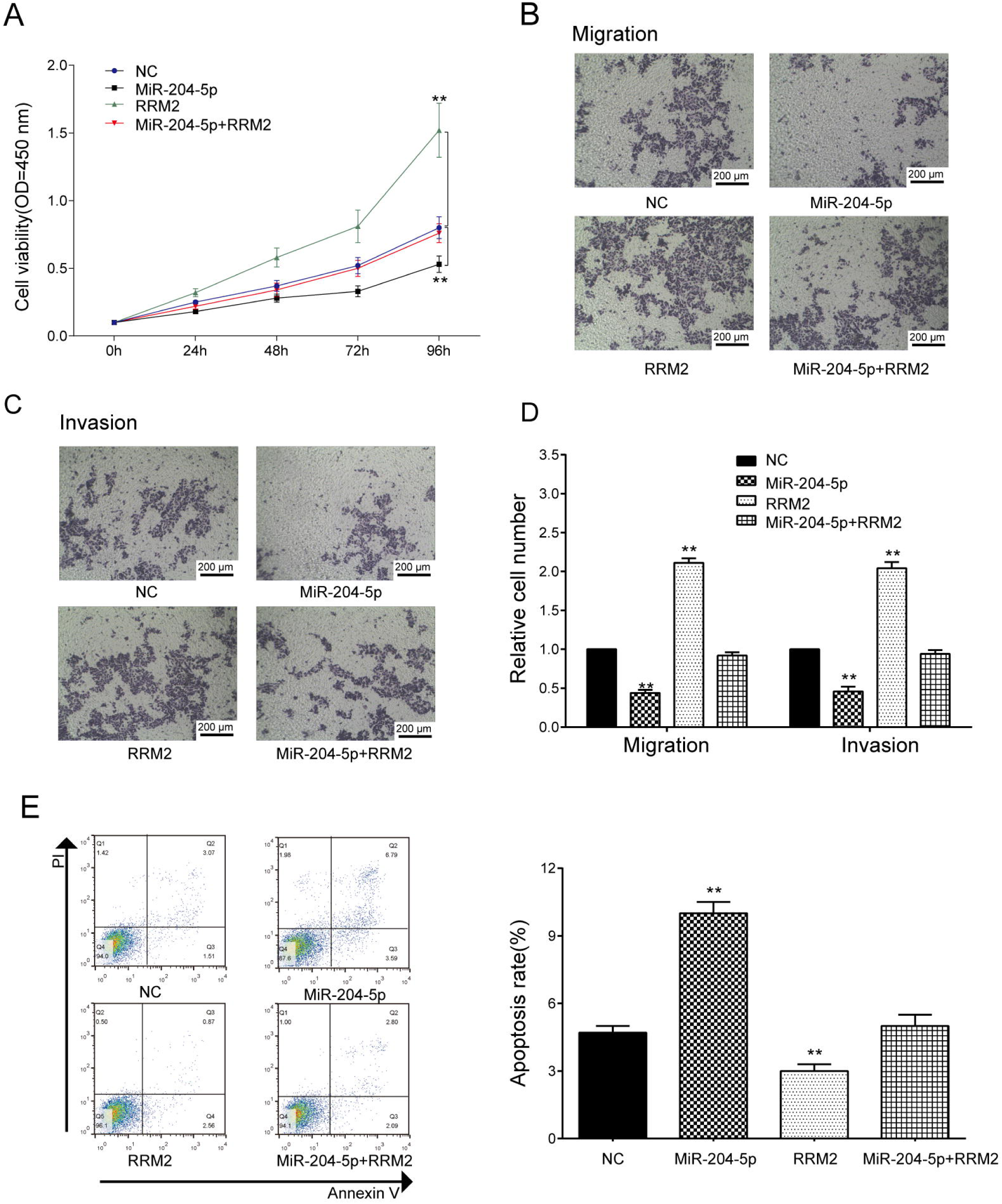
Effect of RRM2 on BC cell activities. (A) CCK-8 assay indicated that cell vitality of HCC1937 cells transfected with RRM2 was higher than that cells transfected with NC. (B-D) Transwell assay showed that compared with NC group, more migrating or more invasive cells were found in RRM2 group while less metastasis cells were observed in miR-204-5p group. (E) Flow cytometry indicated that apoptosis rate in RRM2 group was lower than that in NC group, while apoptosis rate in miR-204-5p group was higher than that in NC group. ^**^ *P*<0.01, compared with NC group.

### Overexpression of DSCAM-AS1 promoted tumor growth

In order to investigate DSCAM-AS1’s tumor promotion effect in vivo, we separately injected equal numbers (10^6^ or 2 × 10^6^) of negative control (NC), DSCAM-AS1 or sh-DSCAM-AS1 transduced HCC1937 cells into nude mice, the relative expression levels of each group were validated by qRT-PCR. The results of qRT-PCR indicated that DSCAM-AS1 expression was significantly increased in DSCAM-AS1 transfected HCC1937 cell group, while it was decreased in sh-DSCAM-AS1 transfected HCC1937 cell group (Figure 6A, *P*<0.01). Tumor volume was measured every 5 days and tumor weight was detected at day 30. The results showed that though tumor volume and weight were increased in all three groups, while tumor volume were increased significantly in DSCAM-AS1 group compared to other groups, indicating overexpression of DSCAM-AS1 promoted tumor growth and knockdown of DSCAM-AS1 largely promoted tumor growth (Figure 6B-D, *P*<0.05). Besides, qRT-PCR revealed that miR-204-5p expression was down-regulated after overexpression of DSCAM-AS1, whereas the expression of miR-204-5p was up-regulated after knockdown of DSCAM-AS1 in HCC1037 transplants (Figure 6E, *P*<0.01). Furthermore, overexpression of DSCAM-AS1 improved *RRM2* mRNA and protein expressions (Figure 6F-G, *P*<0.01). Therefore, these data lead to the conclusion that DSCAM-AS1 promoted tumor growth by inhibiting miR-204-5p and promoting the expression of *RRM2*.

**Fig. 6.**
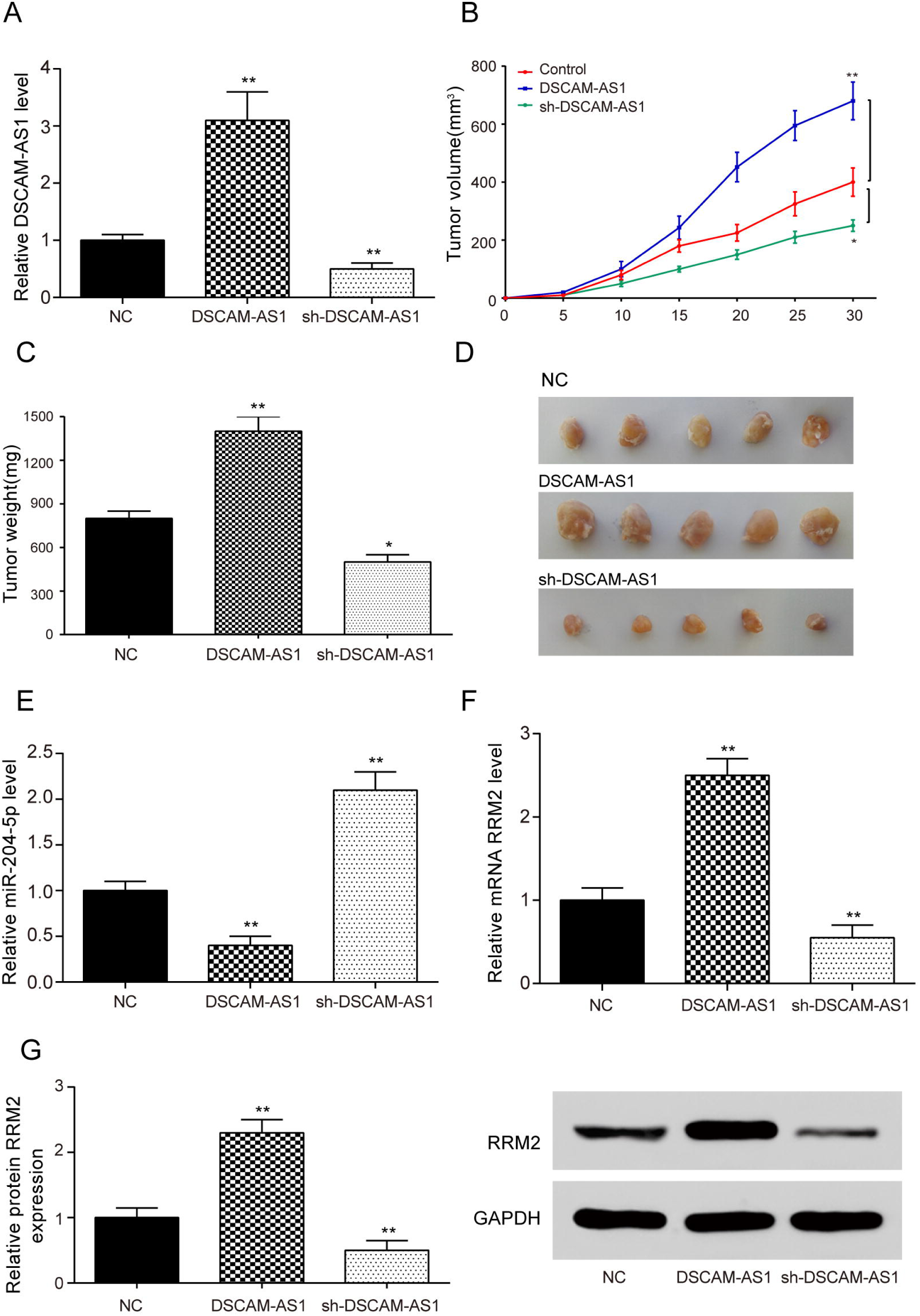
Overexpression of DSCAM-AS1 promoted tumor growth. (A) QRT-PCR indicated that DSCAM-AS1expression of HCC1937 cells transfected with DSCAM-AS1 was significantly higher than that cells transfected with sh-DSCAM-AS1. (B) Tumor volume of DSCAM-AS1 group was larger than that of NC group with the increase of time. (C) Tumor weight was much larger in DSCAM-AS1 group compared with NC group at day 30. (D) Tumor diagram at day 30. (E) QRT-PCR revealed that miR-204-5p expression in DSCAM-AS1 group was significantly lower than that in NC group. (F) QRT-PCR showed that RRM2 expression was significantly higher in DSCAM-AS1 group compared with NC group. (G) Western blot revealed that RRM2 protein expression in DSCAM-AS1 group was significantly higher than that in NC group.

## Discussion

A number of publications have confirmed that lncRNAs are associated with tumorigenesis in BC (Cui et al., 2017). In the present study, the effects of lncRNA DSCAM-AS1 on BC have been investigated. Firstly, we found that DSCAM-AS1 was significantly up-regulated, while miR-204-5p was down-regulated in BC cells through microarray analysis and qRT-PCR. Secondly, we revealed that DSCAM-AS1 directly targeted on miR-204-5p by dual-luciferase reporter system. Meanwhile, the results of CCK-8, transwell and flow cytometry assays indicated that DSCAM-AS1 promoted BC cells proliferation and metastasis and impaired cell apoptosis by impairing the expression of miR-204-5p. Besides, we examined the relationship between miR-204-5p and *RRM2*, and found that miR-204-5p inhibited *RRM2* expression. Moreover, a series of experiments demonstrated that *RRM2* contributed to the promotion of BC cell proliferation and metastasis, as well as the suppression of cell apoptosis. Furthermore, *in vivo* experiments showed that knockdown of DSCAM-AS1 could suppress the tumor growth.

DSCAM-AS1 as an oncogenic lncRNA has been previously reported to be associated with several human cancers (Niknafs et al., 2016). DSCAM-AS1 has been considered as a major discriminant in various cancer cell lines and tumors (Miano et al., 2016). Our experiments revealed that DSCAM-AS1 was up-regulated in BC. Besides, overexpression of DSCAM-AS1 promoted cell reproduction and metastasis as well as inhibited cell apoptosis in BC. A number of similar results have been reported in previous publications. For example, Niknafs *et al.* found that DSCAM-AS1 expressed at a very high level in BC tissues, and promoted the growth and invasion of T47D cells *in vivo* (Niknafs et al., 2016). Miano *et al.* demonstrated that there was an aberrant up-regulation of DSCAM-AS1 in BC samples compared with adjacent tissues, and DSCAM-AS1 was related to cancer cell motility, adhesion and invasion (Miano et al., 2016). Xu *et al.* revealed that the knockdown of DSCAM-AS1 inhibited BC cell proliferation and cycle progression as well as increased cell apoptosis *in vitro* (Xu et al., 2017).

MiR-204-5p has been predicted as an anti-oncogenic molecule in multiple types of cancers (Luo et al., 2017). Several publications have reported that the miR-204-5p expression was down-regulated in various tumors and acted as a potent tumor suppressor inhibiting tumor proliferation and metastasis, including hepatocellular carcinoma, oral squamous cell carcinoma and BC (Jiang et al., 2016; Wang et al., 2016; Zeng et al., 2016). In BC, Li *et al.* showed that the expression of miR-204 significantly reduced compared with adjacent normal BC tissues, and the loss of miR-204 might promote the proliferation and invasive of BC cells (Luo et al., 2017). Additionally, miR-204-5p can interact with lncRNAs to perform an additional regulation of tumor growth. Wang *et al.* found that down-regulation of MALAT1 increased miR-204 expression in BC cells and inhibited cell invasion through reversing epithelial-mesenchymal transition (Wang et al., 2017). Based on previous researches, we further researched the relationship of miR-204-5p and DSCAM-AS1 on BC pathology. The results indicated that miR-204-5p inhibited BC cells reproduction and enhanced cell apoptosis, the expression of which was inhibited by DSCAM-AS1.

*RRM2*, as a key gene in pyrimidine metabolism, was correlated with varied human tumors (Putluri et al., 2014). Previous literatures have reported that *RRM2* had a high expression in cancer and was involved in tumor aggressiveness, poor prognosis, and chemoresistance (Shah et al., 2015). For instance, Sturtz *et al.* demonstrated that *RRM2* expressed at higher levels in tumor tissues, associated with up-regulation of cellular propagation and invasiveness (Sturtz et al., 2014). Shah *et al.* revealed that *RRM2* was overexpressed in tamoxifen-resistant BC cells, and enhanced the proliferation and metastasis of MCF-7 BC cells (Shah et al., 2015). Consistently, we found that *RRM2* was up-regulated in BC. Meanwhile, a series of experimental results indicated that the effects of *RRM2* on BC cell activities were reversely changed by miR-204-5p. We clearly investigated the molecular mechanism underlying DSCAM-AS1, miR-204-5p and *RRM2* and revealed their relationship in BC for the first time.

However, some shortages of our study should be considered in our following researches. For instance, in vitro experiments of DSCAM-AS1 and *RRM2* should be conducted to comprehensively understand the molecular network of DSCAM-AS1, miR-204-5p and *RRM2* in BC.

## Conclusion

In conclusion, we validated that DSCAM-AS1 was elevated in BC cells, DSCAM-AS1 promoted the proliferation and metastasis of BC cells and suppressed cell apoptosis by inhibiting miR-204-5p and increasing *RRM2* expression. Therefore, this study suggests that inhibition of DSCAM-AS1 may be a promising therapeutic strategy for treatment of BC.

## Disclosure

### Competing interests

The authors declare that there are no competing interests associated with the manuscript.

### Ethics approval and consent to participate

All procedures performed in studies involving human participants and animals were in accordance with the ethical standards of the Affiliated Center Hospital, Xinxiang Medical University. Written informed consents were obtained from all individual participants included in the study.

### Consent for publication

The authors consent for publication.

### Availability of data and material

The datasets used and analysed during the current study are available from the corresponding author on reasonable request.

### Authors’ contributions

Substantial contribution to the conception and design of the work: Wen-Hui Liang and Na Li; Analysis and interpretation of the data: Zhi-Qing Yuan, Zhi-Hui Wang and Xin-Lai Qian; Drafting the manuscript: Wen-Hui Liang and Na Li; Revising the work critically for important intellectual content: Na Li; Collecting of grants: Na Li; Final approval of the work: All authors.

### Funding

This study was supported in part by grants from the Natural Science Foundation of Henan Province, China (No.162300410220), the Educational Commission of Henan Province, China (No. 16A310002, No. 18A310004), the Science and Technique Foundation of Henan Province, China (No. 201403133).

## Acknowledgements

None.

## References

Bartonicek, N., Maag, J. L. and Dinger, M. E. (2016). Long noncoding RNAs in cancer: mechanisms of action and technological advancements. Mol Cancer 15, 43.

Cui, H., Zhang, Y., Zhang, Q., Chen, W., Zhao, H. and Liang, J. (2017). A comprehensive genome-wide analysis of long noncoding RNA expression profile in hepatocellular carcinoma. Cancer Med 6, 2932–2941.

Flores-Perez, A., Marchat, L. A., Rodriguez-Cuevas, S., Bautista-Pina, V., Hidalgo-Miranda, A., Ocampo, E. A., Martinez, M. S., Palma-Flores, C., Fonseca-Sanchez, M. A., Astudillo-de la Vega, H. et al. (2016). Dual targeting of ANGPT1 and TGFBR2 genes by miR-204 controls angiogenesis in breast cancer. Sci Rep 6, 34504.

Huarte, M. (2015). The emerging role of lncRNAs in cancer. Nat Med 21, 1253–61.

Imam, J. S., Plyler, J. R., Bansal, H., Prajapati, S., Bansal, S., Rebeles, J., Chen, H. I., Chang, Y. F., Panneerdoss, S., Zoghi, B. et al. (2012). Genomic loss of tumor suppressor miRNA-204 promotes cancer cell migration and invasion by activating AKT/mTOR/Rac1 signaling and actin reorganization. PLoS One 7, e52397.

Iwamoto, K., Nakashiro, K., Tanaka, H., Tokuzen, N. and Hamakawa, H. (2015). Ribonucleotide reductase M2 is a promising molecular target for the treatment of oral squamous cell carcinoma. Int J Oncol 46, 1971–7.

Jiang, G., Wen, L., Zheng, H., Jian, Z. and Deng, W. (2016). miR-204-5p targeting SIRT1 regulates hepatocellular carcinoma progression. Cell Biochem Funct 34, 505–510.

Li, W., Jin, X., Zhang, Q., Zhang, G., Deng, X. and Ma, L. (2014). Decreased expression of miR-204 is associated with poor prognosis in patients with breast cancer. Int J Clin Exp Pathol 7, 3287–92.

Liu, J. and Li, Y. (2015). Trichostatin A and Tamoxifen inhibit breast cancer cell growth by miR-204 and ERalpha reducing AKT/mTOR pathway. Biochem Biophys Res Commun 467, 242–7.

Liu, L., Wang, J., Li, X., Ma, J., Shi, C., Zhu, H., Xi, Q., Zhang, J., Zhao, X. and Gu, M. (2015). MiR-204-5p suppresses cell proliferation by inhibiting IGFBP5 in papillary thyroid carcinoma. Biochem Biophys Res Commun 457, 621–6.

Lu, A. G., Feng, H., Wang, P. X., Han, D. P., Chen, X. H. and Zheng, M. H. (2012). Emerging roles of the ribonucleotide reductase M2 in colorectal cancer and ultraviolet-induced DNA damage repair. World J Gastroenterol 18, 4704–13.

Luo, Y. H., Tang, W., Zhang, X., Tan, Z., Guo, W. L., Zhao, N., Pang, S. M., Dang, Y. W., Rong, M. H. and Cao, J. (2017). Promising significance of the association of miR-204-5p expression with clinicopathological features of hepatocellular carcinoma. Medicine (Baltimore) 96, e7545.

Miano, V., Ferrero, G., Reineri, S., Caizzi, L., Annaratone, L., Ricci, L., Cutrupi, S., Castellano, I., Cordero, F. and De Bortoli, M. (2016). Luminal long non-coding RNAs regulated by estrogen receptor alpha in a ligand-independent manner show functional roles in breast cancer. Oncotarget 7, 3201–16.

Niknafs, Y. S., Han, S., Ma, T., Speers, C., Zhang, C., Wilder-Romans, K., Iyer, M. K., Pitchiaya, S., Malik, R., Hosono, Y. et al. (2016). The lncRNA landscape of breast cancer reveals a role for DSCAM-AS1 in breast cancer progression. Nat Commun 7, 12791.

Putluri, N., Maity, S., Kommagani, R., Creighton, C. J., Putluri, V., Chen, F., Nanda, S., Bhowmik, S. K., Terunuma, A., Dorsey, T. et al. (2014). Pathway-centric integrative analysis identifies RRM2 as a prognostic marker in breast cancer associated with poor survival and tamoxifen resistance. Neoplasia 16, 390–402.

Shah, K. N., Wilson, E. A., Malla, R., Elford, H. L. and Faridi, J. S. (2015). Targeting Ribonucleotide Reductase M2 and NF-kappaB Activation with Didox to Circumvent Tamoxifen Resistance in Breast Cancer. Mol Cancer Ther 14, 2411–21.

Shen, S. Q., Huang, L. S., Xiao, X. L., Zhu, X. F., Xiong, D. D., Cao, X. M., Wei, K. L. Chen, G. and Feng, Z. B. (2017). miR-204 regulates the biological behavior of breast cancer MCF-7 cells by directly targeting FOXA1. Oncol Rep 38, 368–376.

Sturtz, L. A., Deyarmin, B., van Laar, R., Yarina, W., Shriver, C. D. and Ellsworth, R. E. (2014). Gene expression differences in adipose tissue associated with breast tumorigenesis. Adipocyte 3, 107–14.

Vimalraj, S., Miranda, P. J., Ramyakrishna, B. and Selvamurugan,. (2013). Regulation of breast cancer and bone metastasis by microRNAs. Dis Markers 35, 369–87.

Wang, X., Li, F. and Zhou, X. (2016). miR-204-5p regulates cell proliferation and metastasis through inhibiting CXCR4 expression in OSCC. Biomed Pharmacother 82, 202–7.

Wang, X., Qiu, W., Zhang, G., Xu, S., Gao, Q. and Yang, Z. (2015). MicroRNA-204 targets JAK2 in breast cancer and induces cell apoptosis through the STAT3/BCl-2/survivin pathway. Int J Clin Exp Pathol 8, 5017–25.

Wang, Y., Zhou, Y., Yang, Z., Chen, B., Huang, W., Liu, Y. and Zhang, Y. (2017). MiR-204/ZEB2 axis functions as key mediator for MALAT1-induced epithelial-mesenchymal transition in breast cancer. Tumour Biol 39, 1010428317690998.

Xi, J., Feng, J., Li, Q., Li, X. and Zeng, S. (2017). The long non-coding RNA lncFOXO1 suppresses growth of human breast cancer cells through association with BAP1. Int J Oncol 50, 1663–1670.

Xia, Z., Liu, F., Zhang, J. and Liu, L. (2015). Decreased Expression of MiRNA-204-5p Contributes to Glioma Progression and Promotes Glioma Cell Growth, Migration and Invasion. PLoS One 10, e0132399.

Xu, S., Kong, D., Chen, Q., Ping, Y. and Pang, D. (2017). Oncogenic long noncoding RNA landscape in breast cancer. Mol Cancer 16, 129.

Yan, M., Zhang, L., Li, G., Xiao, S., Dai, J. and Cen, X. (2017). Long noncoding RNA linc-ITGB1 promotes cell migration and invasion in human breast cancer. Biotechnol Appl Biochem 64, 5–13.

Zeng, J., Wei, M., Shi, R., Cai, C., Liu, X., Li, T. and Ma, W. (2016). MiR-204-5p/Six1 feedback loop promotes epithelial-mesenchymal transition in breast cancer. Tumour Biol 37, 2729–35.

Zhang, B., Yin, Y., Hu, Y., Zhang, J., Bian, Z., Song, M., Hua, D. and Huang, Z. (2015). MicroRNA-204-5p inhibits gastric cancer cell proliferation by downregulating USP47 and RAB22A. Med Oncol 32, 331.

Zhao, W., Luo, J. and Jiao, S. (2014). Comprehensive characterization of cancer subtype associated long non-coding RNAs and their clinical implications. Sci Rep 4, 6591.

Zhong, Z., Cao, Y., Yang, S. and Zhang, S. (2016). Overexpression of RRM2 in gastric cancer cell promotes their invasiveness via AKT/NF-kappaB signaling pathway. Pharmazie 71, 280–4.

Zhou, W., Ye, X. L., Xu, J., Cao, M. G., Fang, Z. Y., Li, L. Y., Guan, G. H., Liu, Q., Qian, Y. H. and Xie, D. (2017). The lncRNA H19 mediates breast cancer cell plasticity during EMT and MET plasticity by differentially sponging miR-200b/c and let-7b. Sci Signal 10.

